# Novel *Escherichia coli* Phages Representing a Distinct Genus within *Stephanstirmvirinae*: Genome and Host Range Characteristics

**DOI:** 10.1101/2025.02.05.636633

**Authors:** Tomoyoshi Kaneko, Jumpei Uchiyama, Toshifumi Osaka, Satoshi Tsuneda

**Affiliations:** Department of Life Science and Medical Bioscience, Waseda University, Tokyo, Japan; Phage Therapy Institute, Comprehensive Research Organization, Waseda University, Tokyo, Japan; Department of Bacteriology, Graduate School of Medicine Dentistry and Pharmaceutical Sciences, Okayama University; Department of Microbiology and Immunology, Tokyo Women’s Medical University, Tokyo, Japan

**Keywords:** Bacteriophage, *Escherichia coli*, *Stephanstirmvirinae*, *Justusliebigvirus*, *Wecvirus*, Phage taxonomy, Host range, Genomic analysis

## Abstract

Bacteriophages play crucial roles in microbial ecosystems and have potential biotechnological applications. However, our understanding of culturable phages remains limited. This study characterized six novel *Escherichia coli* phages isolated from pig farm wastewater and urban sewage using comprehensive genomic, morphological, and host-range analyses. Using multiple comparative approaches, including gene-sharing network analysis, average nucleotide identity (ANI), and nucleotide intergenomic similarity (NIS), we demonstrated that five of these phages form a distinct group within the subfamily *Stephanstirmvirinae*, potentially representing a novel genus provisionally named "*Wecvirus*”. These phages were further classified into two distinct species within the proposed genus, each of which exhibits a unique host range pattern. This host specificity is reflected in the species-specific differences in the amino acid sequences of tail fibers, which are crucial for infection. The remaining phage, which was not classified as *Wecvirus* exhibited characteristics that challenged the current classification criteria, highlighting the need for more flexible taxonomic approaches. Our findings expand the understanding of phage diversity within *Stephanstirmvirinae* and contribute to the evolving phage taxonomy framework.

**Importance:** The rapid emergence of antibiotic-resistant pathogens necessitates the development of alternative antimicrobial strategies, with phage therapy showing promising potential. However, the successful implementation of phage-based treatments requires a comprehensive understanding of phage diversity and host-pathogen interactions. Our study expands the known diversity within *Stephanstirmvirinae* by characterizing novel *E. coli* phages that constitute a distinct genus. The identification of species-specific host recognition mechanisms within this genus provides insights into phage-host adaptation. These findings contribute to our understanding of phage taxonomy and evolution, while also offering practical implications for the development of phage-based therapeutics against pathogenic *E. coli* strains. This research highlights the importance of continued exploration of novel phages for both fundamental understanding and therapeutic applications.

## Introduction

Bacteriophages (phages) are viruses that infect bacteria and are the most abundant biological entities on Earth (1). They play vital roles in shaping microbial communities, driving bacterial evolution, and maintaining ecological balance in various environments (2–5). The diversity and abundance of phages significantly influences global biogeochemical cycles and holds promise for biotechnological applications, including phage therapy and biocontrol (6–8).

Phage classification is considerably challenging because of rapid evolution and extensive genetic diversity of phages (9). The International Committee on Taxonomy of Viruses (ICTV) has continuously updated its classification system for viruses, including phages, to accommodate new discoveries and genomic data. In 2022, the ICTV substantially revised the classification of *Caudovirales*, abandoning the morphology-based classification in favor of binomial species names (10). This revision has resulted in a classification system that better reflects phylogenetic relationships based on genome sequences.

The subfamily *Stephanstirmvirinae*, a relatively new group within the class *Caudoviricetes*, consists of two genera: *Justusliebigvirus* and *Phapecoctavirus*. Phages in this subfamily primarily infect γ-proteobacteria, including *Escherichia coli*, *Klebsiella pneumoniae*, and members of Pseudomonadales (11–17). Interestingly, *Campylobacter jejuni*, an ε-proteobacterium with distinct outer membrane structure, has also been reported as a host of *Stephanstirmvirinae*, suggesting that phages in this subfamily may possess diverse host recognition mechanisms (18). These bacterial hosts are medically significant because of their rapid development of antimicrobial resistance, and phages infecting these bacteria are considered promising candidates for combating drug-resistant pathogens (6, 7, 19). Furthermore, these phages have been isolated from diverse environments, including water samples (sewage and freshwater), feces (avian and human), and compost, suggesting their ability to infect bacterial strains from different sources (11–13, 18). Phages belonging to *Justusliebigvirus* and *Phapecoctavirus* have been reported to show lytic activity against drug-resistant bacteria, human pathogens, and plant pathogens, leading to ongoing research into their potential as novel antimicrobial agents (13–15, 17). The diversity of host recognition mechanisms in these phages indicates their potential to adapt to different bacterial strains. This characteristic, combined with its demonstrated activity against various pathogenic bacteria, makes it a promising candidate for therapeutic applications. Understanding these host-phage interactions is crucial for developing effective phage-based treatments against emerging drug-resistant bacteria.

As of August 29, 2024, seven publications on “*Phapecoctavirus*” and one on “*Justusliebigvirus*” have been indexed in PubMed (11–18). While these studies have established basic characteristics, such as linear double-stranded DNA genomes (∼150 kbp) and GC content (∼39%), many critical aspects remain unexplored. In particular, the mechanisms underlying host specificity, including the role of structural proteins, such as tail fibers, warrant further investigation to understand their evolutionary adaptations and therapeutic potential.

In this study, we present a comprehensive characterization of six phage strains: four previously isolated from pig farm wastewater (ΦWec179, ΦWec181, ΦWec186, and ΦWec187) and two from urban sewage (ΦWec188 and ΦWec190) (20). Our analysis included a phylogenetic analysis based on genome sequences, morphological features observed using transmission electron microscopy (TEM), and host range evaluation. This study expands our understanding of phage diversity within the *Stephanstirmvirinae* subfamily and contributes significantly to the broader field of phage taxonomy and evolution.

## Results

### Morphological analysis

The TEM images indicated that all the phages were morphologically identical to myoviruses (Fig. 1). The capsids and tails of each phage exhibited similar morphologies.

**Fig. 1.**
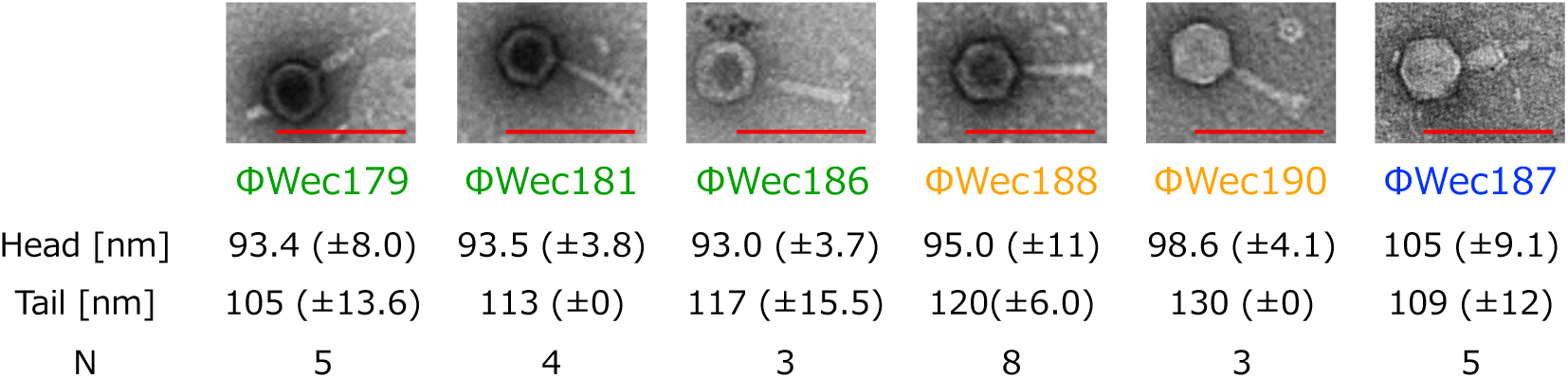
TEM images of phages. Representative TEM images of each phage with mean values and standard deviations for head and tail size. Scale bar represents 200 nm.

### Host range analysis

Host range analysis of multiple *Enterobacteriaceae* species revealed that these phages showed similar patterns, all infecting the screened host, *E. coli* TK001, and *E. coli* clinical isolates, SUTL1, ESBL946, and ESBL991 (Table 1 and Table S1). The ΦWec188 and 190 groups could also infect ESBL1054 and *S. enterica* subsp. *enterica* serovar Pullorum. Furthermore, ΦWec187 could infect all strains susceptible to the other phages plus ESBL1054 and *E. coli* laboratory strains, MG1655 and TOP10F’, and showed lytic activity (without plaque formation) against *E. fergusonii*. The control phages, T4 and T7, infected three laboratory strains (MG1655, TOP10F’, and BL21) and *E. fergusonii*, with T7 also infecting the clinical isolate, ESBL1054.

**Table 1.** Host range of the isolated phages.

| log(EoP) | TK001 | MG1655 | BL21 | TOP10F' | SUTL1 | ESBL946 | ESBL991 | ESBL1054 | IHTL-1 | Sp | *E. fergusonii* |
| --- | --- | --- | --- | --- | --- | --- | --- | --- | --- | --- | --- |
| ΦWec179 | 0.0 | ND | ND | ND | 0.1 | -0.4 | -0.1 | ND | -0.6 | ND | ND |
| ΦWec181 | 0.0 | ND | ND | ND | -0.8 | -0.7 | -1.3 | ND | -0.8 | ND | ND |
| ΦWec186 | 0.0 | ND | ND | ND | -1.2 | -2.1 | -1.4 | ND | -0.8 | ND | ND |
| ΦWec188 | 0.0 | ND | ND | ND | 0.3 | -0.2 | 0.0 | -0.7 | 0.5 | -0.7 | ND |
| ΦWec190 | 0.0 | ND | ND | ND | 0.0 | -0.9 | -0.7 | -0.8 | -0.6 | -0.9 | ND |
| ΦWec187 | 0.0 | 0.9 | ND | 0.5 | 0.8 | 0.1 | -0.1 | 0.2 | 1.4 | ND | pseud |
| T4 | pseud | 0.0 | -0.2 | 0.0 | pseud | pseud | pseud | ND | pseud | pseud | -3.1 |
| T7 | ND | 0.0 | -1.7 | -2.3 | ND | ND | ND | -0.5 | ND | ND | -2.0 |

ND indicates not detected. Sp represents *Salmonella enterica* subsp. *enterica* serovar Pullorum.

### Taxonomic analysis based on whole genome sequences

BLASTn analysis revealed that among the six phages characterized, ΦWec179, 181, and 186 showed highest similarity to *Escherichia* phage vB_EcoM_DE16 (accession no. OP595145.1) of the *Phapecoctavirus* genus within *Stephanstirmvirinae* subfamily, with a similarity (identity × coverage) of 55.32%. ΦWec187, which showed notably lower GC content (37.5%) than other isolated phages (42.2-42.3%), was most closely related to *Escherichia* phage AV123 (accession no. OR352953.1) of the *Justusliebigvirus* genus with 94.17% similarity. ΦWec188 and 190 showed highest similarity to *Escherichia* phage Mt1B1_P17 (accession no. NC_052662.1) of the *Phapecoctavirus* genus, with a similarity of 54.36% (Table S2). The genomic characteristics of the phages are summarized in Table 2.

**Table 2.** Overview of *E. coli* phages ΦWec179, 181, 186, 188, and 190.

| Phage | Genbank accession no. | Genome length bp | GC % | CDS | tRNA | Source |
| --- | --- | --- | --- | --- | --- | --- |
| ΦWec179 | LC739533 | 154,295 | 42.2 | 331 | 10 | pig farm sewage |
| ΦWec181 | LC739534 | 154,402 | 42.2 | 331 | 10 | pig farm sewage |
| ΦWec186 | LC739535 | 154,402 | 42.2 | 331 | 10 | pig farm sewage |
| ΦWec187 | LC739536 | 145,321 | 37.5 | 275 | 15 | pig farm sewage |
| ΦWec188 | LC739537 | 149,683 | 42.3 | 325 | 11 | urban sewage |
| ΦWec190 | LC739539 | 150,541 | 42.3 | 328 | 11 | urban sewage |

To more thoroughly investigate the relationships between these isolated phages, we conducted a gene-sharing network analysis based on shared protein clusters between viral genomes using vContact2 (Fig. 2A, Table S3). ΦWec179, 181, 186, 188, and 190 formed their own unique cluster, VC82_1 (Fig. 2B, Table S3). ΦWec187 belonged to VC82_0, which contains only one reference phage, *Enterobacteria* phage phi92 (Fig. 2B, Table S3). The phylogenetically related cluster, VC82_2, contained five reference phages: *Escherichia* phage ESCO13, *Escherichia* phage ESCO5, *Escherichia* phage phAPEC8, *Escherichia* phage vB_EcoM_Schickermooser, and *Klebsiella* phage ZCKP1. All reference phages in VC82_0 and VC82_2 belonged to the subfamily *Stephanstirmvirinae*. To evaluate the similarity between all NCBI-registered *Stephanstirmvirinae* subfamily phages (as of July 24, 2024) and ΦWec179, 181, 186, 187, 188, and 190, we calculated the nucleotide intergenomic similarity (NIS) using VIRIDIC and average nucleotide identity (ANI) using pyani (Fig. 3, Table S4). ΦWec187 showed NIS values below 95.5% when compared with the registered *Stephanstirmvirinae* phages, whereas its ANI values were between 0.7% and 94%. Notably, phages with accession numbers MG065645.1, MG065689.1, MG065639.1, MG065656.1, OX090893.1, and LR865361.1, showed NIS values between 95% and 95.5% with ΦWec187, and their ANI values were approximately 90%. However, some phages with ANI values above 90% had NIS values below 95%, whereas others with ANI values below 90% (approximately 89%) had NIS values above 95%. ΦWec179, 181, 186, 188, and 190 showed NIS values below 66% and ANI values between 20-50% when compared with registered *Stephanstirmvirinae* phages. Furthermore, both NIS and ANI values for ΦWec179, 181, 186, 188, and 190 exceeded 70%; they formed two distinct groups, the ΦWec179, 181, and 186 and ΦWec188 and 190 groups, when a 90% threshold was applied.

**Fig. 2.**
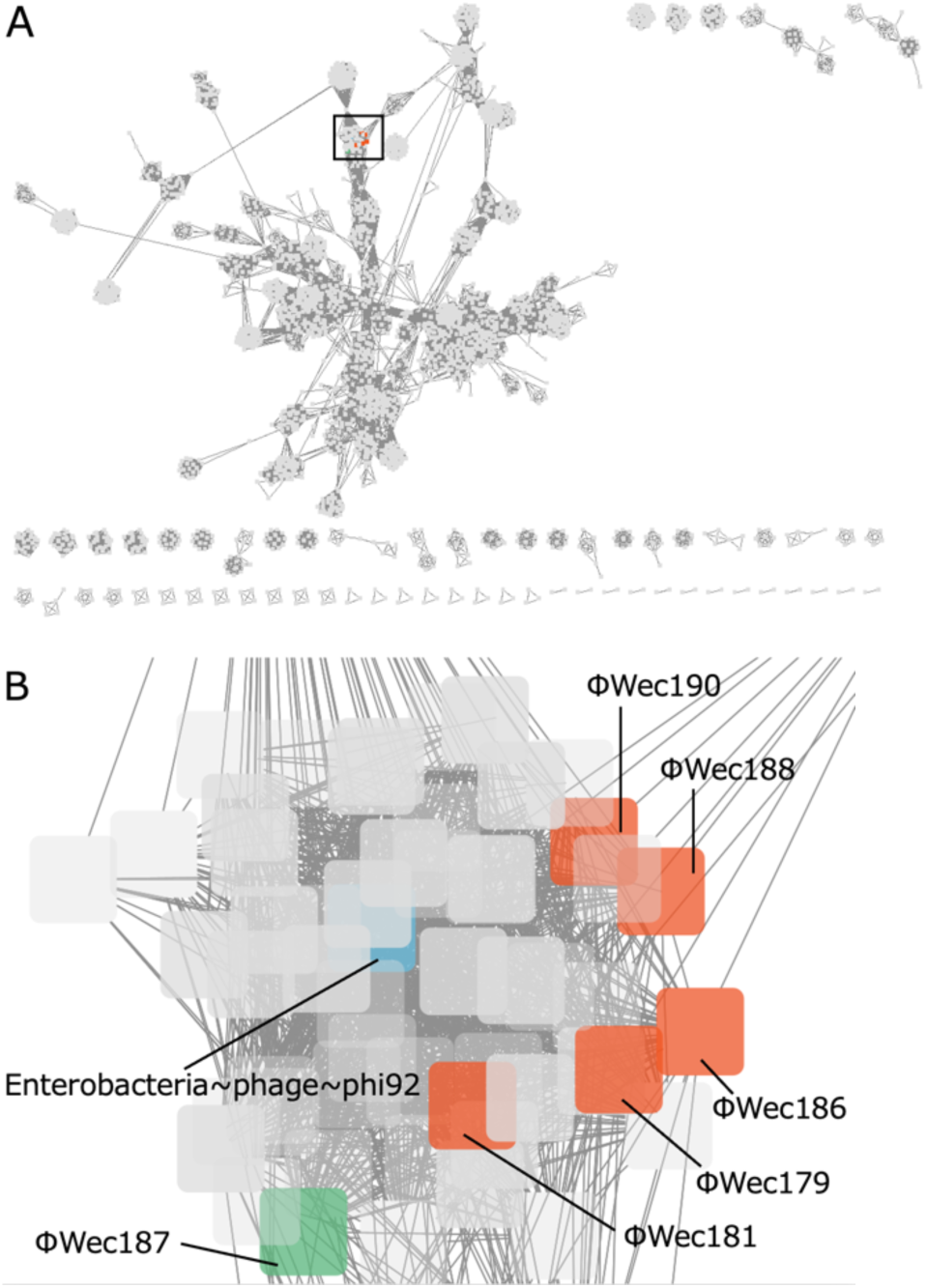
Gene-sharing network analysis of phage genomes. (A) Network visualization of gene-sharing relationships among 3,509 phage genomes. The boxed area indicates the VC82 cluster. (B) Enlarged view of the VC82 cluster. Orange nodes indicate ΦWec179, 181, 186, 188, and 190 (VC82_1), green node indicates ΦWec187 (VC82_0), light blue node indicates the single VC82 reference phage, and gray nodes indicate other reference phages.

**Fig. 3.**
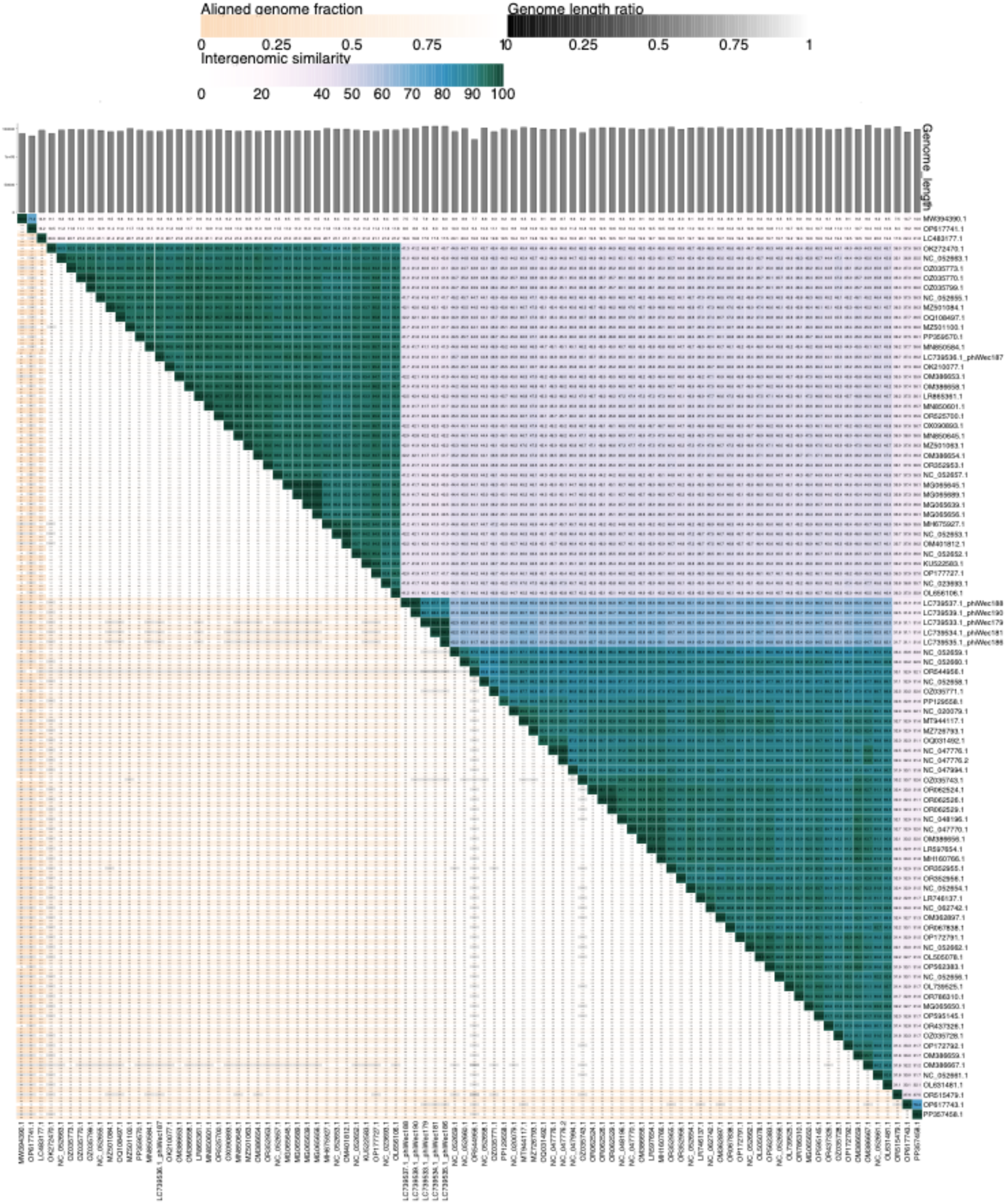
Similarity analysis of phage isolates of the subfamily *Stephanstirmvirinae*. NIS, genome length, and aligned genome fractions were determined using VIRIDIC. The phage genome information for comparison was obtained from registered *Stephanstirmvirinae* using the genbank_get_genomes_by_taxon.py script included in the pyani package.

Comparative genomic analysis visually confirmed the high similarity suggested by NIS and ANI values within the ΦWec179, 181, and 186 and ΦWec188 and 190 groups (Fig. 4A, B). Comparison between ΦWec179 and ΦWec188 revealed some similarity, although it was less pronounced than that within groups (Fig. 4C). Comparison between representative strains ΦWec179 (from the ΦWec179, 181, and 186 group), ΦWec188 (from the ΦWec188, 190 group), and ΦWec187 showed negligible similarity.

**Fig. 4.**
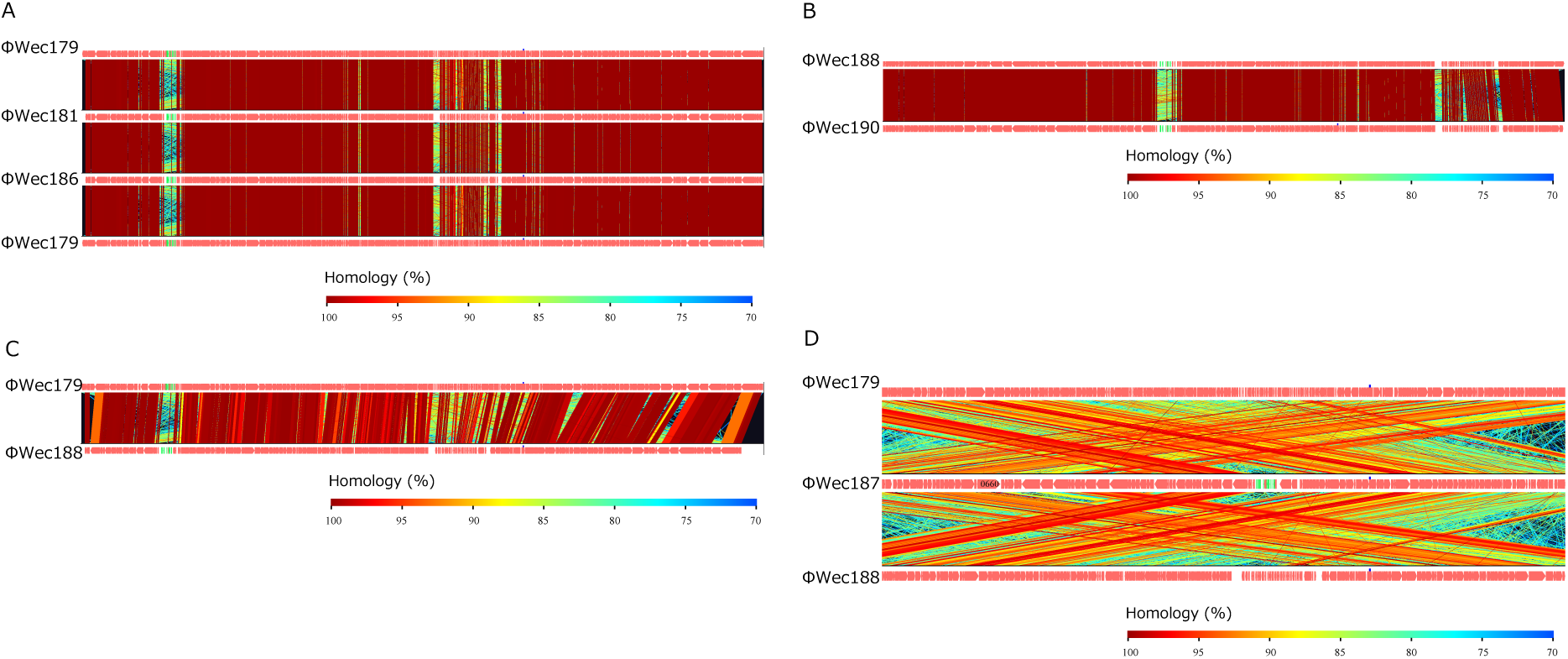
Comparative genomic analysis of isolated phages. Heatmap representation of genome similarities. (A) ΦWec179, 181, and 186 group. (B) ΦWec188 and 190. (C) ΦWec179 and 188. (D) ΦWec179, 187, and 188 group.

### Phylogenetic analysis of tail fiber proteins

Six types of tail fiber protein-coding genes were identified based on homology with the VOG database using the results of Pharokka annotation: TfpL (long-tail fiber protein; 943 or 958 amino acids or AAs), TfpM1 (major tail fiber protein-1; 890 or 891 AAs), TfpM2 (major tail fiber protein-2; 681 AAs), TfpS1 (short-tail fiber protein-1; 353 AAs), TfpS2 (short-tail fiber protein-2; 235 AAs), and TfpS3 (short-tail fiber protein-3; 157 AAs). These genes were present in all phages except ΦWec187, which lacked the gene encoding TfpM1.

Molecular phylogenetic analysis revealed that for all analyzed tail fiber genes, the ΦWec179, 181, and 186 group and ΦWec188 and 190 groups formed distinct clusters (Fig. 5A-F). The phylogenetic position of ΦWec187 varied depending on the gene; TfpM2, TfpS1, TfpS2, and TfpS3 showed independent positions in both groups (Fig. 5C-F). However, in the TfpL phylogenetic tree, ΦWec187 was included in the same clade as the ΦWec188 group, although it maintained an independent position within this group (Fig. 5A).

**Fig. 5.**
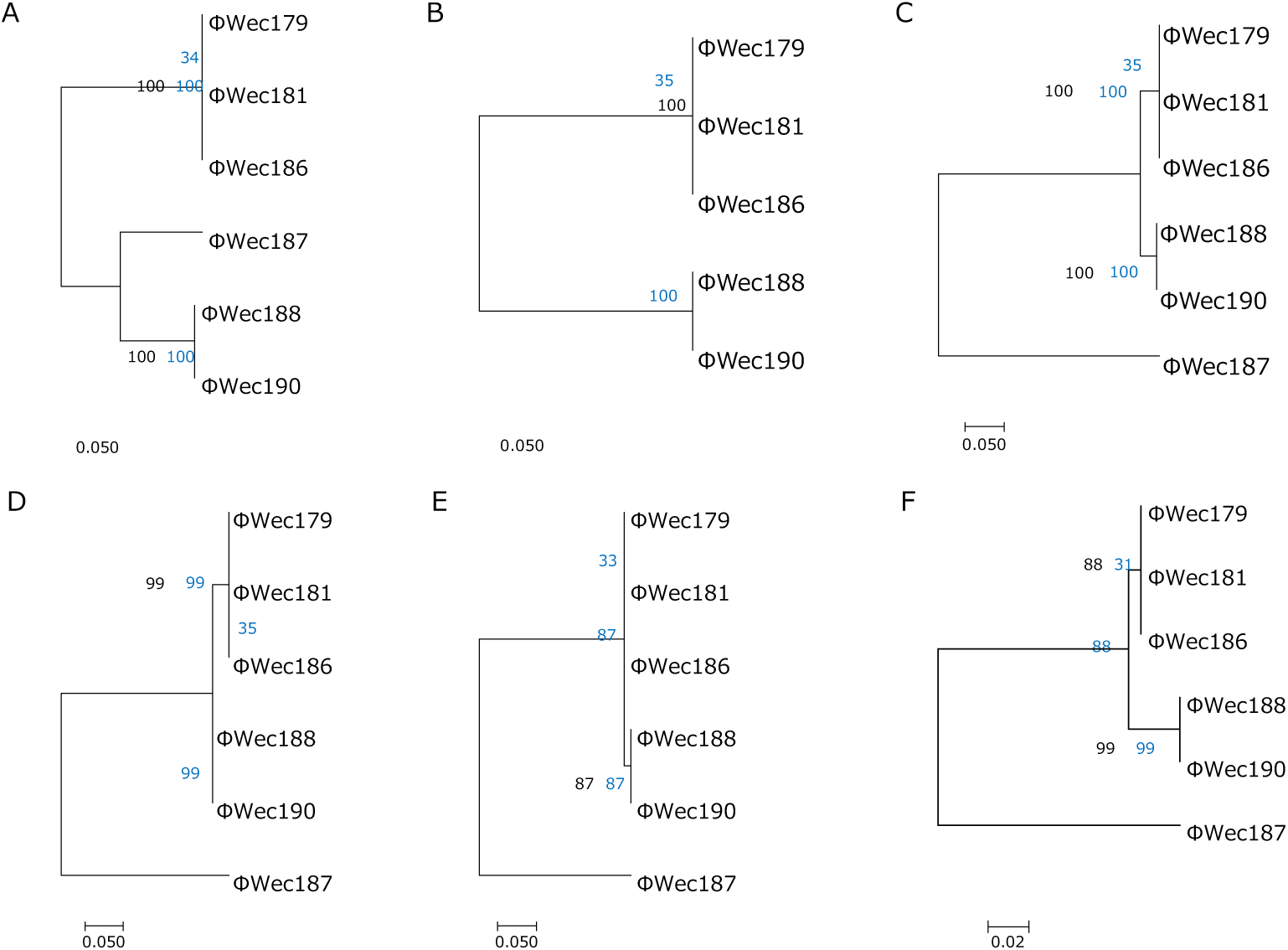
Molecular phylogenetic analysis of tail fiber genes. Molecular phylogenetic trees based on the amino acid sequences of tail fiber genes. Trees were constructed using the ML method with a bootstrap value of 1,000. Black text shows the original tree and blue text shows the bootstrap values. (A) TfpL; (B) TfpM1; (C) TfpM2; (D) TfpS1; (E) TfpS2; (F) TfpS3.

Furthermore, while each tail fiber protein-coding gene (TfpL, TfpM1, TfpM2, TfpS1, TfpS2, and TfpS3) was completely conserved within the ΦWec179 group and ΦWec188 group, slight differences were observed between the groups, with variations concentrated in the C-terminal regions. Notably, although these genes were perfectly conserved within the ΦWec188 group, their order was not conserved. Additionally, the arrangement of conserved tail fiber genes (TfpL, TfpM2, TfpS1, TfpS2, and TfpS3) in ΦWec187 matched that in ΦWec188.

## Discussion

In this study, we characterized six novel *E. coli* phages using genomic, morphological, and host range analyses. Our findings revealed that five of these phages (ΦWec179, ΦWec181, ΦWec186, ΦWec188, and ΦWec190) represent a new genus within *Stephanstirmvirinae*, which we propose naming "*Wecvirus*”. These phages form two distinct species, based on their genomic similarities and host range patterns. The remaining phage, ΦWec187, shows characteristics of *Justusliebigvirus* although it cannot be classified accurately under the current classification criteria.

Morphological analysis confirmed that all the isolated phages belonged to the myovirus morphotype, consistent with that observed for other members of *Stephanstirmvirinae* (12–14, 16). Host range patterns revealed distinct infection profiles in the isolated phages. From the perspective of host range testing, all the environmental phages analyzed in this study, including T4 (*Caudoviricetes*; *Straboviridae*; *Tevenvirinae*) and T7 (*Autographiviridae*; *Studiervirinae*; *Teseptimavirus*), which are phylogenetically distinct from *Wecvirus*, showed approximately 13% infectivity, confirming the narrow host range typical of *Enterobacteriaceae*-infecting phages (21). The differential ability to infect laboratory strains versus clinical isolates suggests an adaptation to specific host populations. Notably, the broader host range of ΦWec187 compared to those of other isolates indicates potential differences in host recognition mechanisms. Our findings align with those of previous studies on *Stephanstirmvirinae* members, where similar variations in host specificity were reported. For instance, Nicolas *et al*. reported that *Phapecoctavirus* phages showed approximately 24% infectivity against a panel of 32 *E. coli* strains, primarily avian pathogenic *E. coli* (APEC) strains, whereas *Wecvirus* isolates demonstrated a narrower host range against different clinical isolates (12). Similarly, Markuskovál *et al*. observed that the *Justusliebigvirus* phage had a 23% infection rate against uropathogenic *E. coli* (17). These differences in host specificity possibly reflect the diverse origins of the tested bacterial strains rather than inherent differences in host range breadth.

Taxonomic analysis revealed significant evolutionary relationships within *Stephanstirmvirinae*. The ΦWec179, ΦWec181, ΦWec186, ΦWec188, and ΦWec190 phages formed a distinct cluster and showed similarity values below the 70% genus-level threshold in both NIS and ANI analyses when compared with known members. This discovery expands our understanding of the diversity within *Stephanstirmvirinae*, suggesting that the current sampling has not yet captured the full range of phage diversity within this subfamily.

Among these five phages, NIS and ANI analyses revealed two distinct clusters with high internal similarities (>95%). Between-group similarity values of 80-88% indicated that these clusters represent separate species, despite their close evolutionary relationship. The consistently higher NIS values (60-65%), compared to the corresponding ANI values (20-50%), between these phages and reference phages reflect fundamental differences in how these metrics are calculated. NIS, which performs direct whole-genome comparisons, appears to capture conserved regions and overall genome structure more effectively. Conversely, ANI values, which average similarities across fragmented sequences (22), may better evaluate local similarities but do not completely account for phage-specific genomic features, such as extensive rearrangements or large-scale insertions/deletions.

We observed discrepancies between the NIS and ANI values of ΦWec187, making classification of this phage under the current taxonomic criteria challenging. The unique characteristics of this phage highlight the complexity of current phage classification systems and warrant further investigation. We propose that ΦWec187 should be temporarily classified as “unclassified *Stephanstirmvirinae*” and investigated in the future.

Tail fibers are composed of multiple proteins that play crucial roles in host recognition (23, 24). For example, the proximal and distal tail fibers of the T4 phage have distinct protein compositions (25). The multiple tail fiber genes identified in this study possibly constitute different parts of the tail structure and may enable the recognition of different host molecules (26).

Detailed analysis of tail fiber proteins provides insights into phage-host interactions and adaptation mechanisms. The differences in host ranges correlate with group-specific characteristics of tail fiber composition, particularly in the C-terminal regions, which are crucial for receptor recognition (27–30). While whole-genome comparisons showed clear grouping patterns, tail fiber phylogenetic analysis revealed more complex relationships, particularly regarding ΦWec187, suggesting mosaic evolution via horizontal gene transfer rather than simple linear evolution (9, 31).

Our previous study demonstrated that all five phages target the R1 core lipopolysaccharide (LPS), with both groups specifically recognizing galactose transferred by WaaT (32). The C-terminal sequence variations in tail fibers suggest the acquisition of additional host recognition capabilities while maintaining this core LPS-binding function. In particular, the ΦWec179-186 group appears to utilize flagella as a secondary receptor, while the ΦWec188-190 group requires the inner membrane protein, YhaH, indicating distinct evolutionary trajectories in host adaptation.

Subtle differences in the host range were observed, even within the groups. For example, ΦWec179, 181, and 186, despite belonging to the same group and showing complete conservation of tail fiber protein sequences, exhibited slight variations in infection efficiency against certain strains. These observations suggest that while tail structure largely determines host range, host utilization efficiency may involve factors beyond phage adsorption, such as replication efficiency within host cells. For instance, the T4 collar and whiskers regulate long-tail fiber retraction under unfavorable conditions (33), and bacterial phage defense systems can reduce efficiency of plating (EoP) (34). Future research should focus on the functional analysis of receptor-binding proteins and investigation of host cell surface interactions. Additionally, the elucidation of genes with unknown functions may contribute to our understanding of host specificity.

Based on these results and per the ICTV naming conventions (Rules 3.21, 3.22) (35), we propose the following taxonomic classifications:

1. Establishment of a new genus, “*Wecvirus*”, within the subfamily *Stephanstirmvirinae*
2. Designation of a new species, “*Wecvirus wec179*”, comprising ΦWec179, ΦWec181, and ΦWec186
3. Designation of a new species, “*Wecvirus wec188*”, comprising ΦWec188 and ΦWec190

These names follow the ICTV guidelines (35), with the genus name consisting of a single word with the suffix “-virus” and the species names comprising two words: the genus name followed by the isolate number. The new genus name is based on the representative isolate designation (Wec). The *Wecvirus* genus is genetically distinct from known *Stephanstirmvirinae* subfamily members (ANI and NIS <70%), and each of its new species “*Wecvirus wec179*” and “*Wecvirus wec188*”, display unique host ranges. The establishment of this new genus and its constituent species enhances our understanding of phage diversity and refines the taxonomic system within the *Stephanstirmvirinae* subfamily. Future research is expected to identify additional phages belonging to the *Wecvirus* genus and elucidate its ecological role through functional analysis of receptor- binding proteins and investigation of host-phage interactions.

## Materials and methods

### Reagents and bacteria

Luria-Bertani medium was used for bacterial and phage cultures. The concentration of soft agar was 0.5%. The *E. coli* strain TK001 used in this study was isolated from the feces of mouse with dextran sodium sulfate (DSS)-induced colitis and was used as the host strain for phage screening (20). The animal experiments for generating mice with DSS-induced colitis used for the isolation of TK001 were reviewed and approved by the Waseda University Academic Research Ethics Committee (approval number: 2020- A009).

### Phage preparation

Four *E. coli* phages (ΦWec179, 181, 186, and 187) were isolated from swine farm effluent and two phages (ΦWec188 and 190) were isolated from municipal sewage. These six phages (ΦWec179, ΦWec181, ΦWec186, ΦWec187, ΦWec188, and ΦWec190) were propagated using *E. coli* TK001 as the host, while T4 and T7 phages were propagated using MG1655 as the host, all using the plate lysate method. Briefly, an overnight bacterial culture (100 μL) and phage suspension (10^4^ PFU/mL, 100 μL) were added to soft agar (0.5%) and overlaid onto agar plates. After overnight incubation (37°C), 5 mL of sodium chloride-magnesium sulfate (SM) buffer was added to the plates, followed by orbital shaking (approximately 250 rpm, 2-3 h). The supernatant was collected in 15 mL tubes and centrifuged (7,300 G, 15 min, 4°C; TOMY MX-305 centrifuge with an AR500- 03 rotor) to separate the solid and liquid components. The liquid component was transferred to new 15 mL tubes, followed by the addition of 1 mL of chloroform. After vortexing and settling, the preparation was used as the phage stock. T4 and T7 were obtained from the Biological Resource Center of the National Institute of Technology and Evaluation in Kisarazu, Japan (NBRC 20004 and 20005, respectively).

### Host range analysis

Phage infectivity against bacterial strains was evaluated as follows: Each phage (approximately 10^7^ PFU/mL) was serially diluted tenfold up to 10^7^ dilution in SM buffer, 1 μL of which was spotted. The EoP was calculated by dividing the titer against each bacterial strain by the titer against the reference host (TK001 for ΦWec179, 181, 186, 188, and 190; MG1655 for T4 and T7). The bacterial strains tested included 33 laboratory and clinical isolates of *E. coli*, four different serotypes of *Salmonella enterica* (NBRC3163, 3313, 13245, and 15335), and one strain of *E. fergusonii* (NBRC102419), details of which are provided in Table S5.

### TEM

Phages (10^9^-10^10^ PFU/mL) were purified via polyethylene glycol precipitation and loaded onto copper grids using a support film (cat. no. 649; Nissin EM Co. Ltd., Tokyo, Japan). After washing thrice with distilled water, the samples were stained with four-fold diluted EM stainer (cat. no. 336; Nissin EM Co., Ltd., Tokyo, Japan) for 1 min. The excess solution was removed with filter paper, and the grids were air-dried before observation using a transmission electron microscope (100 kV, JEM-1010, JEOL, Tokyo, Japan).

### Taxonomic analysis based on whole genome sequences

Previously published genome sequences (LC739533–LC739537 and LC739539) were used (20). Gene-sharing network analysis was conducted using vConTACT2 with reference genomes of 3,503 phages (ProkaryoticViralRefSeq201) and visualized using Cytoscape v3.10.1 (36)(37). Taxonomic information that was not updated within vConTACT2 was manually updated using a taxonomy browser (https://www.ncbi.nlm.nih.gov/Taxonomy/Browser/wwwtax.cgi). ANI analysis was performed using pyani with the "anib" parameter (38). NIS was calculated using VIRIDIC (39). The phage genomes used for the ANI and NIS comparisons were obtained using the Genbank_get_genomes_by_taxon. py script included in the Pyani package. Comparative genome analysis was performed using Genomematcher in blastp mode (40).

### Genome annotation

Phage annotations were performed using Pharokka to complement previous annotations conducted using RAST and DFAST (41). A phylogenetic tree was constructed using MEGA11 with the maximum likelihood method and 1,000 bootstrap replicates (42). Provisional naming conventions were used to facilitate comparison of the tail fiber protein-coding genes. Based on the amino acid residue counts, the genes were designated as follows: TfpL (943 or 958 residues), TfpM1 (890 or 891 residues), TfpM2 (681 residues), TfpS1 (353 residues), TfpS2 (235 residues), and TfpS3 (157 residues). These designations are used for discussion purposes in this study and are not proposed as official gene names.

## Authorship contributions

Tomoyoshi Kaneko: Conceptualization, Data curation, formal analysis, writing–original draft, investigation

Jumpei Uchiyama: Supervision, Writing - review & editing

Osaka Toshifumi: Supervision, Writing - review & editing

Satoshi Tsuneda: Supervision, Writing - review & editing

## Acknowledgements

This research received no specific grant from any funding agency in the public, commercial, or not-for-profit sectors. The initial English translation of this manuscript was facilitated using the DeepL Translator (DeepL GmbH), ChatGPT-3.5 (OpenAI), Claude 3.5 Sonnet (Anthropic), and Gemini 1.5 Flash (Google DeepMind). We thank Editage (www.editage.com) for professional English language editing. We carefully reviewed and verified the accuracy of the content at each stage of translation and editing.

